# Fawn bedsite selection by a large ungulate living in a peri-urban area

**DOI:** 10.1101/2023.08.11.552922

**Authors:** Kimberly Conteddu, Katie Wilson, Bawan Amin, Srivats Chari, Amy Haigh, Laura L. Griffin, Matthew Quinn, Andrew Ryan, Adam F. Smith, Simone Ciuti

## Abstract

Human-wildlife conflict in expanding peri-urban and urban areas is of increasing concern, as a result of growing human populations along with the associated anthropogenic footprint on wildlife habitats. Empirical data from wildlife research carried out within human dominated landscapes are key to understanding the effects of human pressures on wildlife ecology and behaviour, exploring wildlife behavioural flexibility (or phenotypic plasticity), and informing wildlife management decisions. Here, we explored how female fallow deer (*Dama dama*) responded to human and dog presence during the birthing period in the largest walled urban park in Europe. We collected data on 477 bedsites utilised by 283 neonate fawns across three consecutive fawning seasons, gathered fine-scale data on humans and dogs space use, and built Resource Selection Functions at multiple spatial scales. We found that, when choosing bedsites to give birth and leave fawns unattended, fallow deer mothers significantly avoided hotspots of park visitors on foot (and their dogs) along the hiking trail routes. Bedsites were also unlikely to be in close proximity of paved roads used by vehicle traffic. Additionally, fallow deer mothers were found to select for dense understory vegetation for bedsites, providing low visibility to conceal their offspring. Our results provide detailed insights into bedsite spatial and habitat selection by a large herbivore in response to human activities, and we provide clear indications to wildlife managers to preserve established fawning sites and alleviate human-wildlife conflict during a critical period of the deer annual biological cycle.

## Introduction

We are currently in the Anthropocene, defined by extensive human impact on our planet’s ecology and geology (Morelli et al., 2020). It is estimated that more than 50% of the human population will be concentrated in cities by 2030 (Lowry et al., 2013, Shochat et al., 2006), with these cities presently expanding at an unprecedented rate worldwide (Wolff et al., 2018). This process leads to increased synurbanisation, i.e. wildlife increasingly living in and adapting to urban environments, with a concurrent increase in encounters between humans and wildlife (Barber et al., 2010, Gallo et al., 2019, Huijbers et al., 2013, Tucker et al., 2018). Several concerns have been raised about the potential for artificial selection processes to occur as a result of human-wildlife interactions (i.e., certain morphological and/or behavioural traits favoured over others) within urban settings (Griffin et al., 2022, Lowry et al., 2013).

Empirical research has repeatedly shown that human presence can affect wildlife behaviour significantly (Gander and Ingold, 1997, Hebblewhite et al., 2005, Recarte et al., 1998, Shen-Jin et al., 2007, Whittington et al., 2004). Wildlife species may increase flight response in locations with high human presence (Recarte et al., 1998), as well as increase selection of safer habitats and reduced movement (Gander and Ingold, 1997, Hebblewhite et al., 2005, Recarte et al., 1998, Shen-Jin et al., 2007, Whittington et al., 2004). Within urban settings, however, wildlife species need to adapt their anti-predator strategies (e.g., reduced flight distance) as fleeing any time that they encounter humans would be energetically expensive. This highlights how behavioural flexibility (or phenotypic plasticity) may be an important characteristic for succeeding in urban environments (Lowry et al., 2013).

Dog owners within urban areas need to walk their dogs and make any possible use of green areas and urban parks, potentially leading to high encounter rate between dogs and wildlife inhabiting urban areas. Dog presence has been shown to alter wildlife behaviour, though responses vary depending on the wildlife species (Manor and Saltz, 2004, Martinetto and Cugnasse, 2001, Miller et al., 2001). Both domestic and feral dog presence increases alert and escape distances in many ungulate species, and has been associated with a reduction in offspring:female ratios (Manor and Saltz, 2004, Martinetto and Cugnasse, 2001, Miller et al., 2001). These decreases in offspring production could be caused by associated increases in vigilance in mothers, decreased breeding behaviour, or a combination of both (St. Clair and Forrest, 2009). Due to the importance of successful breeding seasons in keeping wild animal populations healthy, changes in breeding behaviours during this sensitive period can lead to indirect effects, with potential long-term consequences for the composition (i.e., age ratios) of wild populations and natural selection processes (Lowry et al., 2013). As humans and their dogs are both likely to come into more regular contact with wild animals within increasingly human dominated environments, studies assessing the consequences of human and dog presence on wildlife are increasingly important.

Ungulates exhibit various strategies to increase offspring survival during their first weeks of life; this constitutes a continuum ranging from “hider” to “follower” species (Walther, 1961a). As a “hider” species, fallow deer (*Dama dama*) mothers hide their offspring in dense vegetation, where they remain separated for several hours at a time (Chapman and Chapman, 1997). Selection of these “bedsites” after parturition plays a major role in increasing offspring survival (Gerlach and Vaughan, 1991, Tull et al., 2001, Van Moorter et al., 2009, Monestier et al., 2015). During this vulnerable time for offspring, fallow deer mothers decrease their home-range sizes and select for denser vegetation to increase concealment and decrease predators’ ability to detect fawns (Ciuti et al., 2006). However, research to date has focused almost exclusively on rural areas, with little understanding of bedsite selection in peri-urban landscapes (Smith and Lecount, 1979, Long et al., 1998, Piccolo et al., 2010). This highlights the need for an empirical quantification of the impact of people and their dogs on deer behaviour during fawning to manage wildlife populations within urban settings.

Here, we gathered data on 477 bedsites used by 283 individual fallow deer fawns across 3 consecutive fawning seasons in the largest walled urban park in Europe, Phoenix Park, located at the heart of Dublin, Ireland. This is the largest dataset ever compiled on bedsite selection in a large ungulate in an urban parkland. We explored bedsite selection by fallow deer mothers that have adapted to live in a peri-urban area under high human pressure. Phoenix Park is an ideal model for this study as boasts a substantial herd of wild fallow deer, it receives 10 million annual visitors, typically on foot with many walking their dogs, and large amounts of vehicular traffic (which includes commuters to and from Dublin city centre) at the edge of a capital city. Also, because of the walled nature of the park and the peri-urban environment, it is extremely hard for the deer to leave and find suitable habitat outside the park.

We made 3 main predictions related to our hypothesis that fallow deer mothers would select for the most suitable bedsite features while attempting to minimise human disturbance, where possible, in such a restricted space: (i) mothers would select bedsites characterized by denser vegetation and lower visibility, when compared to random available sites, to reduce fawn detectability by terrestrial predators (including humans); (ii) bedsites would be located outside and at greater distances from hotspots of human and dog presence along hiking trails; and (iii) mothers would avoid roads with vehicle traffic when selecting fawns’ bedsites. Our study site, Phoenix park, is dominated by park visitors during the day; as a consequence, fallow deer mothers are expected to find the best balance between selecting suitable vegetation and an appropriate distance from humans for their fawns’ bedsites, giving us the opportunity to describe deer plasticity and adaptation to extreme conditions in terms of human presence. To better disentangle such behavioural response, we analysed our data looking at two spatial scales: large scale (study-site-level), to depict general avoidance of human presence hotspots; and a small scale (bedsite-level), to understand specific decisions made by mothers when selecting for bedsite characteristics and visibility. Our work has important management implications because our analytical approach allows to estimate threshold distances (distance to roads and hiking trails) and vegetation characteristics (vegetation density and related visibility) that may be used by park managers to create buffer zones and strictly protected core areas for wild deer during this important stage of their annual biological cycle.

## Materials and methods

### Study site

The study was conducted in Phoenix Park, a 709ha urban park located in Dublin, Ireland (53°22’N, -6°21’W, Fig. 1), which receives an estimated 10 million visitors annually (Office of Public Works, OPW, official data). The vegetation is mainly grassland (56%) and woodland (31%), with 7% of the park covered by roads and buildings (OPW, 2020). The Park management (OPW) maintains a fallow deer population of around 600 individuals (end of summer population estimate, which includes newborn fawns, OPW official data), and this number is kept constant by yearly culls. The only natural predator in the park is the red fox, which seldomly predates newborn fawns. Over the years, however, we have made several direct observations of unleashed dogs chasing fawns, in many cases resulting in the injury or death of fawns.

**Fig. 1:**
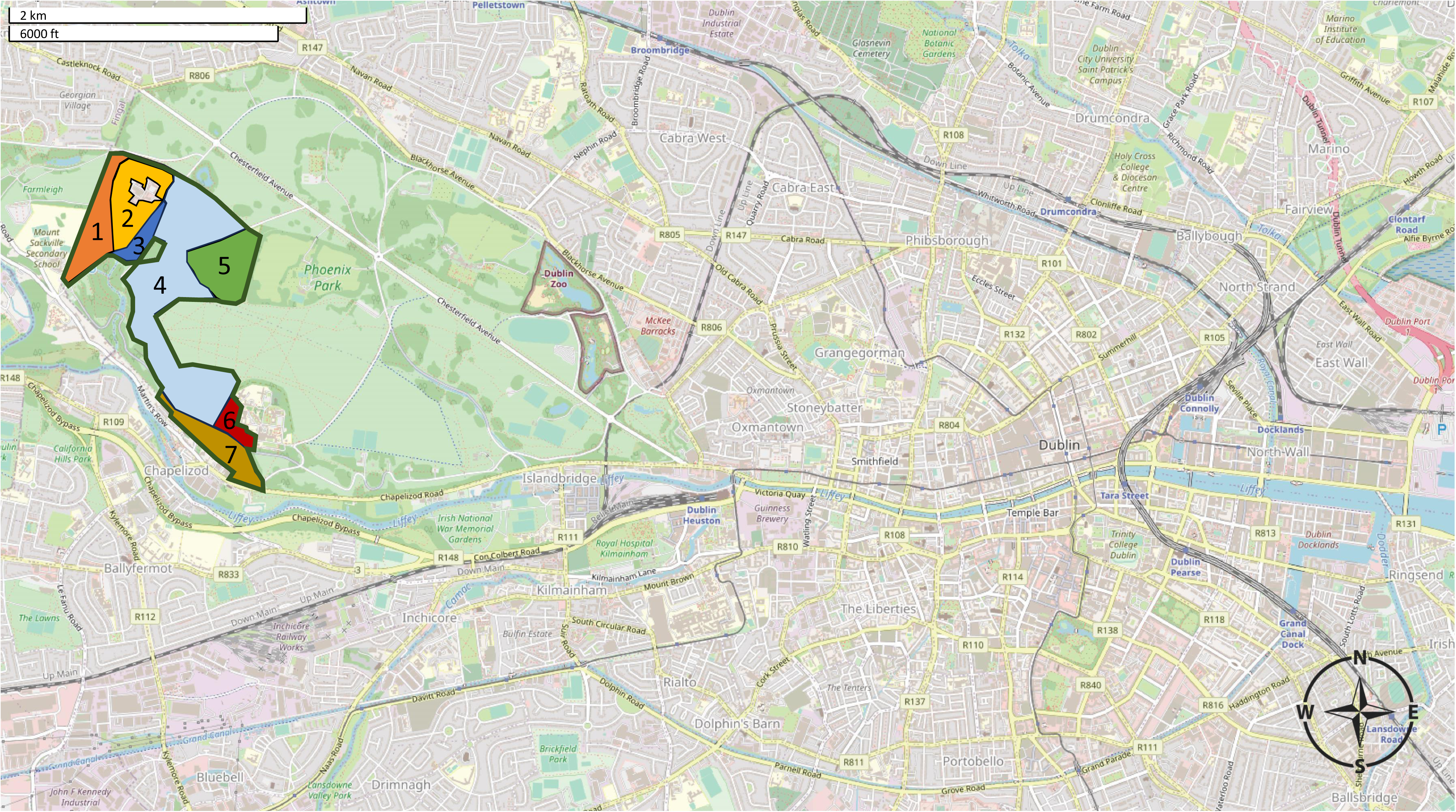
Map of the Dublin metropolitan area, with the study site location, Phoenix Park, clearly visible as the largest green space in the top left of the map. The highlighted polygon (in green) in the western side of Phoenix Park represents the fallow deer fawning area, with the 7 sectors clearly defined.

### Data collection

#### Fawn captures and bedsite characteristics

Fawns have routinely been captured in June (from June 5^th^ to June 28^th^) and ear-tagged with unique numbered plastic tags (Allflex medium; Mullinahone Co-op, Ireland) since the early 1970s as part of the monitoring and management of the fallow deer herd (Hayden et al., 1992, Amin et al., 2021). We split the geographical area typically used by females as fawning sites (displayed in both Fig 1 and Fig. 2a) into seven sectors. These were patrolled routinely, visiting 2-3 sectors daily, with each sector visited at least once every 2-3 days. Approximately 10-15 trained team members supervised by a certified wildlife biologist captured fawns using circular fishing nets (1–1.5m of diameter) manoeuvred by long handles (1.5m long). The capture team usually worked between 9:00 to 16:00 hrs daily, 7 days a week for more than three consecutive weeks, with the goal of tagging most of the newborn fawns within this population (∼85%).

**Fig. 2:**
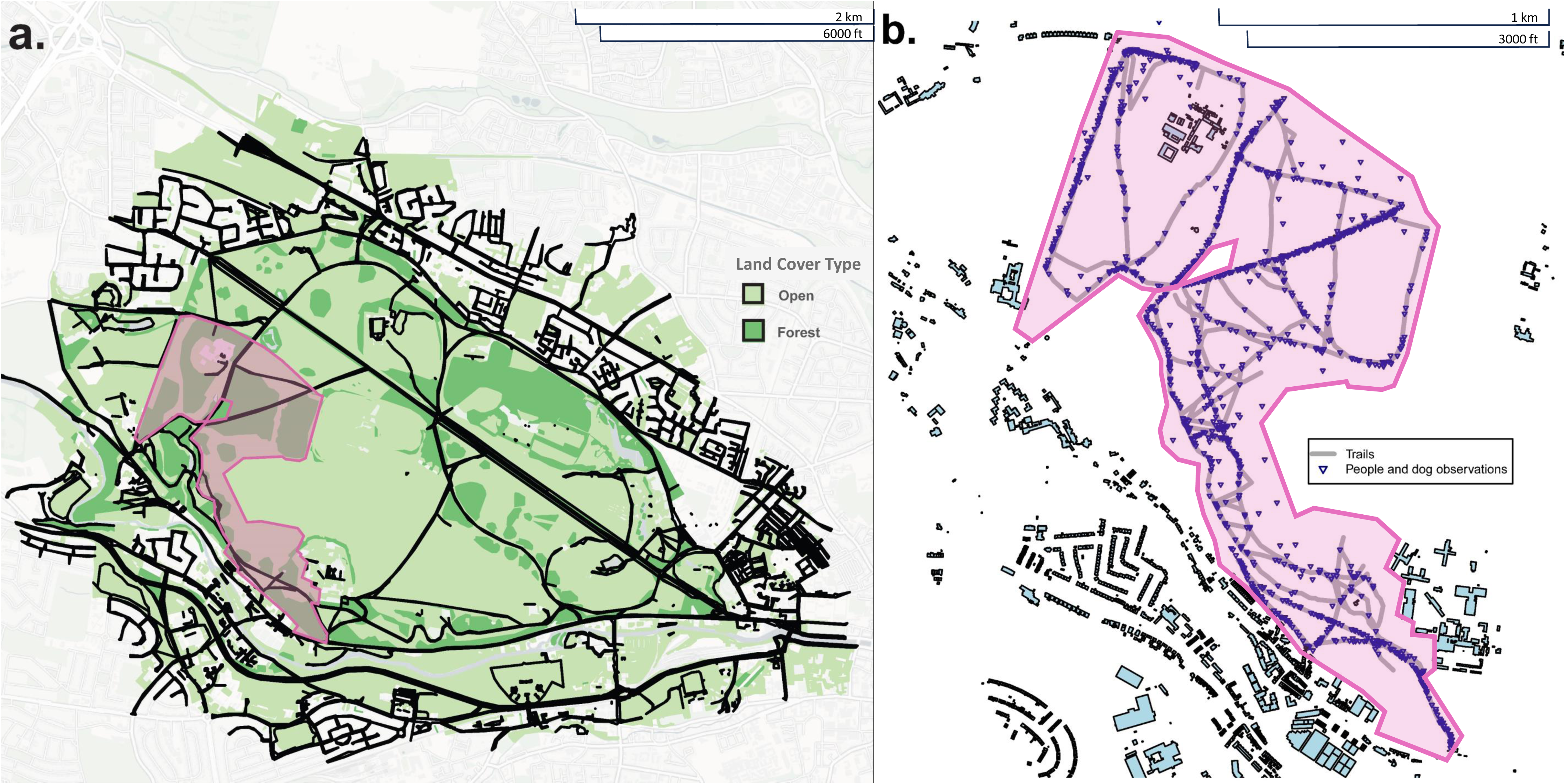
Map of the Phoenix Park, Dublin (a) including land cover types (open, forest), roads (black linear features. Within the fawning area, (b) hiking trails (grey lines) were systematically walked during the fawning season to locate park visitors and their dogs (blue triangles). The fallow deer fawning area, which has been patrolled by the team during ear-tagging procedures, is indicated by the pink polygon in both plots.

The capture team collected data on fawns and their associated bedsites following a well-defined protocol; see Amin et al. (2021) for a full description of the information collected on fawns. For this study, we used the following data: fawn identity (ear tag code) and sex; fawn weight (i.e. an objective proxy for age *sensu* Amin et al. (2021), determined by placing the fawn into a 100-litre bag, which was weighed using a digital scale – resolution: 0.01 kg, Dario Markenartikelvertrieb); GPS coordinates of the bedsite (using a Garmin etrex 30 hand-held unit); and bedsite visibility (see below for full description). Fawns were routinely recaptured to monitor body growth (Griffin et al., 2023), allowing us to collect bedsite characteristics in older fawns. Our final database consisted of fawn capture data collected in 2018 (n = 102 individual fawns), 2019 (n = 81), and 2020 (n = 100). Note that we also had data on 4 additional bedsites outside the capture sectors which we excluded due to the lack of random available points. Because of recaptures, we gathered data on a total of 477 bedsites (n = 163 in 2018, n = 152 in 2019, and n = 162 in 2020, Fig. 3). Note that the bedsite location of recaptured fawns could be different from the one of the first capture, either because the mother moved the fawn, the fawn – once old enough to move independently (1-2 weeks old) – decided to relocate elsewhere, or because of our disturbance. Nevertheless, we were interested in capturing the behavioural process behind the selection of bedsites in an area heavily used and disturbed by humans, and therefore keen to learn from multiple recaptures of the same individual, with the *a priori* assumption that when a bedsite was disturbed, the next one would be selected to minimise human disturbance. Concurrent research in the same park (Faull et al. 2024) has showed that bedsite selection is dictated by the mother when the fawns are neonates (first few days of life). Older (heavier) fawns, however, tend to be more mobile and may have a role in selecting the exact place where to settle down when moving from one bedsite to another (Faull et al. 2024).

**Fig. 3:**
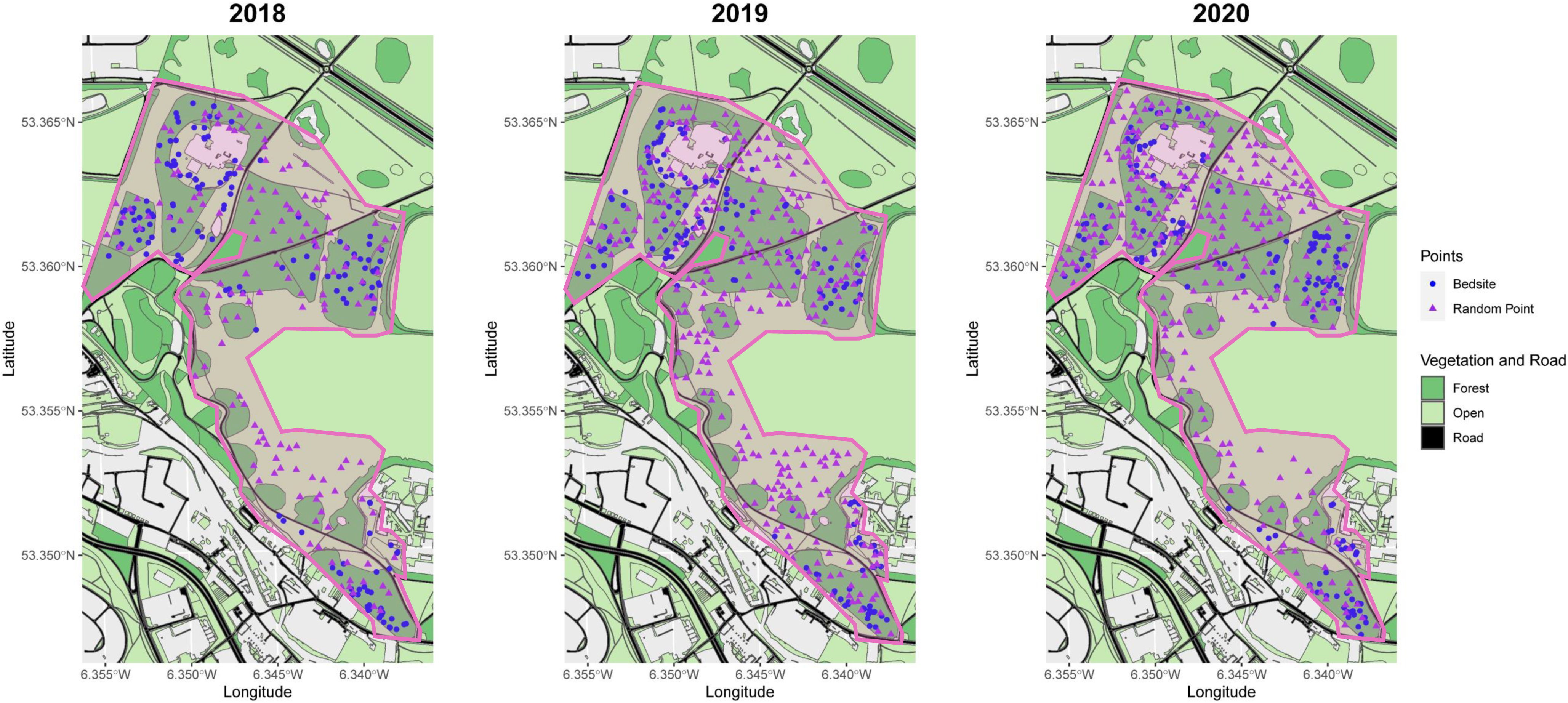
Locations of fallow deer fawn bedsites (blue circle) and random available points (purple triangle) across the 3 years of the study (2018-2020) within the traditional fawning area (pink area) of the Phoenix Park, Dublin.

We assessed the average visibility of the fawn bedsite – ranging from 0 (no visibility) to 1 (full visibility) (*sensu* Bongi et al., 2008) – as follows: from a distance of 10m along the four cardinal directions we recorded the proportion of visually unobstructed triangles on a cardboard shape the size of a standing fawn (height: 45cm, width: 36cm), with the bedsite set as the centroid. We observed the cardboard shape from an eye height of 70cm, which approximately corresponds to the eye height of a red fox or a medium-sized dog. The study protocol and all research procedures were approved by the UCD Animal Research Ethics Committee for the duration of 5 years (2018-2023), under the permit AREC-E-18-28.

#### Presence-available study design

We used a presence-availability design to depict bedsite selection using Resource Selection Functions (RSFs) (Thurfjell et al., 2014, Ciuti et al., 2018). RSFs allowed us to estimate the relative probability of selection by comparing environmental features of used sites (a.k.a. presence data from sites where fawns were found and captured) to those of sites selected at random, defining environmental characteristics available in the study site. Random available points were selected in R 3.6.3 (R Core Team, 2020) within the capture areas with the rule that they each had to be at least 20m apart to avoid overlap (and therefore dependency) of visibility measurements taken at each bedsite. Random points were visited during the fawning season, and, for each location, we recorded the visibility following the same protocol adopted for the true fawn bedsites (Fig. 3).

#### Spatially-explicit data collection on park visitors and their dogs during the fawning season

We collected data on park visitors and their dogs during the fawning season of 2019 (7^th^-28^th^ of June, Fig. 2b) to build a spatially-explicit model of human abundance. Hiking trails were the same in 2018, 2019, 2020, and our goal was not to estimate the variation in human abundance across years, but rather understand which hiking trails were commonly used by park visitors. We walked at a constant pace (5km/hr) along all hiking trails following a strict systematic stratification, to survey each trail equally as a function of time of the day and day of the week. We used a GPS handheld unit to record the position of human and dog presence on paths or trails. If people or dogs were outside the path or trail (see Fig. 2b) – which occurred seldomly – their GPS location was estimated by combining data from the GPS hand-held unit (Garmin etrex 30), a rangefinder (Hawke Endurance), and a compass. For each observation, we counted the number of visitors and dogs.

### Data Analysis

All analyses were carried out in R 3.6.3 (R Core Team, 2020). Using the human and dog presence observations, we created probability maps of park visitors and dogs in the fawning area. We first overlaid a grid of 50×50m cells on the fawning area and added the observation counts to each cell, once for humans and once for dogs, and separately for weekdays and the weekend, resulting in four observation maps. Since the observations of dogs were considerably lower than the observations of humans, dog presence was converted into a binomial value (presence-absence). We then fitted two spatial Generalized Additive Models (GAMs; Wood, 2017) for humans and dogs as a function of the fawning area location (east coordinate, north coordinate) and day of the week (categorical, 2 levels: Mon-Fri weekday, and Sat-Sun weekend), specified in R as follows: (i) Human abundance model: *gam*(human abundance ∼ s(east coordinate, north coordinate, k=29) + day of the week, family = Poisson (quasi-Poisson corrected)); (ii) Dog presence model: *gam*(dog presence ∼ s(east coordinate, north coordinate, k=29) + day of the week, family = binomial). We used a quasi-Poisson-corrected Poisson error distribution for the human abundance model to account for a mild overdispersion (Zuur et al., 2009). For both models, the spline knots were manually set to 29 to avoid overfitting the data following the indications from the *gam.check* function (Wood, 2017). Both models were used to predict human abundance (number of visitors) and dog presence at the 50×50m resolution using the *predict* function in *mgcv* (Wood, 2017).

For each used and available point we assigned land cover type (Ordinance Survey Ireland, official data – 10m resolution) using the o*ver* function from the *sp* package (i.e. categorical variable with two levels: forest, and open habitat – corresponding to grassland), as well as predicted (either weekday or weekend) human abundance and dog presence, and distance to the nearest paved road (where vehicular traffic was allowed, in meters) using the function *gDistance* from the *sp* package (Pebesma and Bivand, 2005, Bivand et al., 2013). We created buffers of 250m and 2500m around all bedsites to assign the random available points at the two spatial scales (bedsite-level and study-site-level). The small size diameter was chosen since it was the smallest buffer that allowed for all bedsites to be compared with enough available points at this resolution (used:available ratio 1:6). The large scale corresponded to the distance between the two most distant vertices of the polygon defining the fawning area, therefore in this instance availability was defined at the population level (Manly et al. 2007). Buffer zones were created using the *gBuffer* function from the *sp* package in R. Our sampling design did not match presence data with available points in the close proximity only, which should have been analysed using a conditional logistic regression able to stratify on matching pairs where each stratum would have its own intercept (*sensu* Thurfjell et al. 2014). We were interested instead in building multiple scale resource selection functions, meaning that given a spatial scale – e.g. 250m – the environmental characteristics of used points were compared to those of the random available points occurring within the 250 meters of all used points. This allowed us to understand the decisions taken by mothers when selecting for fawn bedsites at different spatial scales: when looking at 250m scale, for instance, we were investigating the selection of environmental characteristics on a small scale, but unable to understand the effect of larger scale patterns (e.g. the avoidance of human hotspots at a greater distance than 250 meters) that could be depicted instead by the large scale analysis.

We screened all variables described above, to include in the RSF, for collinearity (Dormann et al., 2013), and found that the spatially-explicit predictions of number of park visitors were highly collinear with dog presence (see Supporting information). We therefore retained the number of park visitors since it was continuous data and able to give a better picture of different levels of disturbances when compared to the binomial probability of dog presence.

Resource Selection Functions, built assuming an exponential form, require normally distributed continuous predictors (Thurfjell et al., 2014). Therefore, we transformed both human abundance and road distance data to meet model assumptions. We succeeded with the transformation of human abundance data using a box-cox transformation, a family of power transformations which selects the best mathematical function for the data transformation to achieve data normality (i.e., achieved using a log-likelihood procedure to find lambda and transform the non-normally distributed variable, see Supporting information) using the *MASS* package (Venables and Ripley, 2002). We were not able to improve the extremely skewed distribution of road distance; this variable was therefore converted into a two-level categorical predictor (close vs far) using a threshold point of 50m. The threshold point was selected by running a sensitivity analysis with preliminary resource selection functions fitted with different threshold values, and the best model was the one with the lowest Akaike Information Criterion, AIC (see Supporting information).

To estimate the coefficients for the small and large scale RSFs, we fitted Generalized Mixed-effect Models (GLMM, *glmer* function from package *lme4* - Bates et al., 2015) with a binomial distribution of errors (response variable: used points, 1s; and random available points, 0s) and fawn identity as random intercept. The *a priori* model structure – fitted for both spatial scales - is reported in Table 1. We scaled numerical predictors to improve model convergence and included quadratic terms to account for non-linear effects. We defined our model structure *a priori* by considering the predictors that we hypothesized to be ecologically relevant in driving habitat selection in this population living in a peri-urban area. We therefore intentionally did not attempt to simplify the model structure, but rather opted for interpreting the full model structure (Dorman et al. 2018). The inclusion of quadratic effects was motivated by the fact that we did not expect bedsite visibility, distance to roads, nor human abundance to have a straight linear effect but rather a plateau effect (e.g. flat after a certain value of the predictor variable) or fully quadratic effect. Note that sex, weight of the fawn (proxy for age), and year of study were included in the model as confounding predictors to specify the dependency structure of the data (see Table 1 for full details) but no ecological inference will be made because their significant levels would simply predict that e.g. more used locations have been collected in a year compared to the next one. Sex and year could have been included as random intercepts in the GLMM but this was avoided due to reduced degrees of freedom.

**Table 1:** Structure of the GLMM models (binomial distribution of errors) fitted to estimate the parameters for the Resource Selection Functions at two spatial scales (250 and 2500m, respectively) and explaining fawn bedsite selection in the fallow deer population of Phoenix Park, Dublin, Ireland.

| Variable type | Variable name | *A priori* rationale for including the variable in the model |
| --- | --- | --- |
| Response variable | Used, available (1,0) | Response variable. |
| Explanatory (inclusive of quadratic term) | Bedsite visibility (ranging from 0 to 1) | Bedsites were expected to be characterised by lower visibility. Relationship was expected to be non-linear *a priori*. |
| Explanatory variable | Road distance (lower or equal to 50m, greater than 50m) | Bedsites were expected to be less likely located close to roads. |
| Explanatory variable (inclusive of quadratic term) | Human abundance (predicted number of people per 50x50m pixel) | Bedsites were expected to be selected in areas with reduced probability of encountering humans and their dogs (being dogs presence highly correlated with human abundance). Relationship was expected to be non-linear *a priori*. |
| Explanatory variable | Land cover type (2 levels: forest, open) | Bedsites were expected to be selected in habitats with greater cover, i.e. forest. |
| Confounding variable | Year (3 levels: 2018, 2019, 2020) | To specify to the model the dependency structure of the data, with data collected either in 2018, 2019, or 2020. Note that this was not included as random effect due to low degrees of freedom. This variable is included in the model as a confounding but should not be interpreted ecologically. |
| Confounding variable (inclusive of quadratic term) | Fawn weight (in kg) | To account for age of the fawn (older fawns = heavier) and specify to the model the dependency structure of the data. This variable is included in the model as a confounding but should not be interpreted ecologically because not included as interaction term. |
| Confounding variable | Sex of fawn (male, female) | To account for sex of the and specify to the model the dependency structure of the data<br>Note that this was not included as random effect due to low degrees of freedom. This variable is included in the model as a confounding but should not be interpreted ecologically. |
| Random effect | Ear-Tag (identification code of the fawn) | To account for repeated observations of multiple besites belonging to the same fawn. |

Parameters estimated by the GLMM were extracted and plugged into the RSF taking the exponential form (Thurfjell et al., 2014, Fechter et al., 2019, Brivio et al., 2019). RSF scenarios estimating relative probabilities of selection (Thurfjell et al., 2014) were built by varying the predictor of interest (e.g., visibility) while keeping all the other variables to median values (for numerical variables) or to a fixed level (for categorical variables: Year = 2018, Sex = f, Land cover type = open).

## Results

We depicted human abundance and dog presence probability maps (50×50m resolution) of the fawning area in Phoenix Park as predicted by the Generalized Additive Models in Fig. 4 (see Supporting Information for full details). In terms of bedsite use, the n = 477 bedsites were characterised by a mean visibility of 0.06 (min-max: 0-1); 21% of them were at a distance lower than 50m from roads (average distance from roads 119m; min-max: 4-338m); mean human abundance predicted at bedsites was 1.22 people per 50×50m pixels(min-max: 0.0009-5.81), and 70% of the besites were found in forest habitat.

**Fig. 4:**
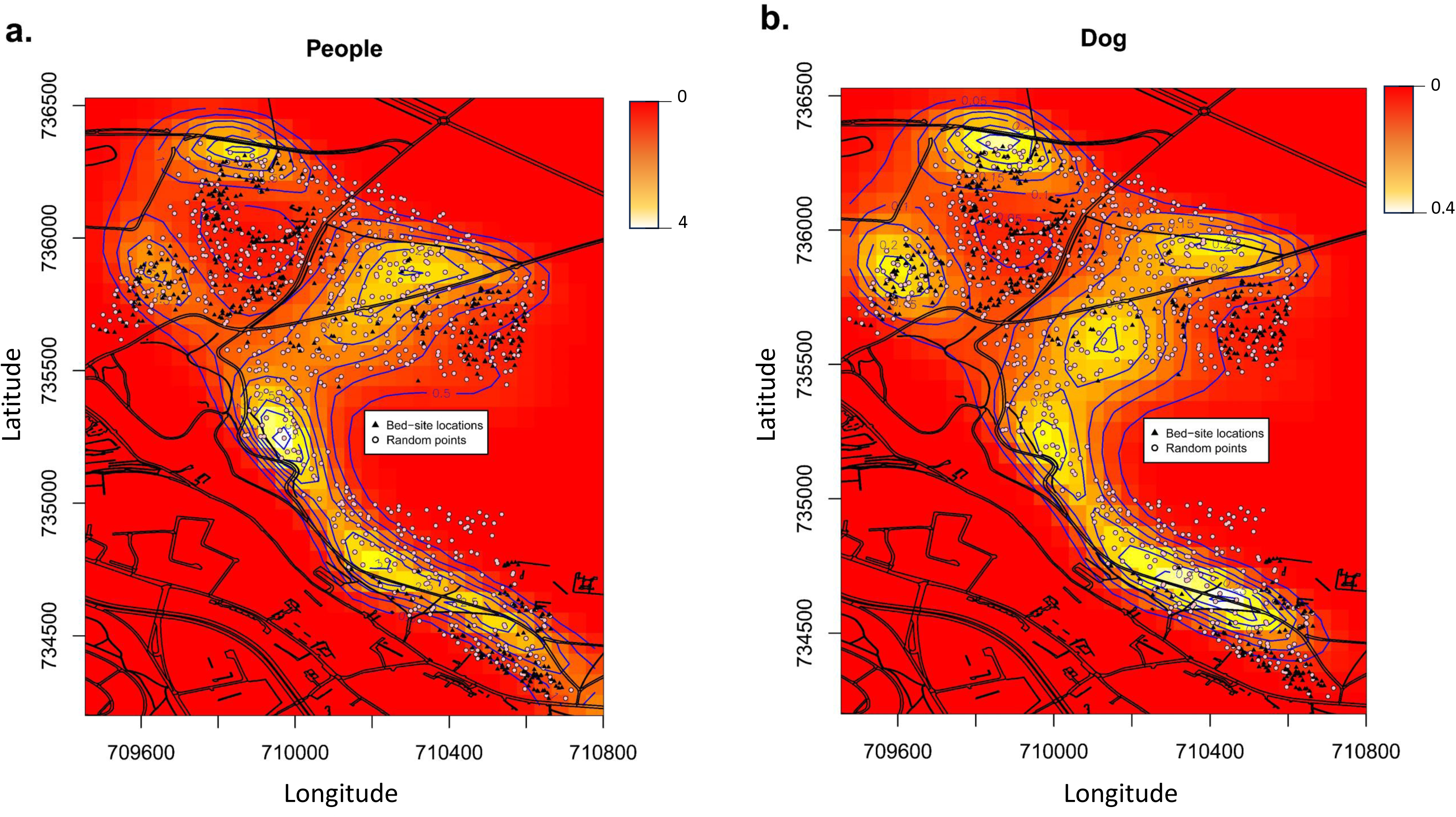
Predicted maps depicting human abundance (a: greater abundance indicated by lighter colours, see legend indicating predicted human abundance) and dog presence (b: greater probability indicated by lighter colours, see legend indicating predicted human presence) in the fawning area of Phoenix Park. Both maps include bedsite locations (triangle) and random available points (circle). X-axis (corresponding to longitude) and Y-axis (corresponding to latitude) are expressed in meters. Note how the location of the fawn bedsites are outside the hotspots of people and dog occurrence.

We reported the parameter estimates of the two GLMMs (small and large scale, respectively) in Table 2. Year, sex, and weight were included in the model to specify the dependency structure of the data, but no inference was made because not included as interaction terms *a priori*. Areas with higher human abundance were avoided at the large spatial scale (Table 2), with bedsites clustered and located outside human hotspots (Fig. 4), whereas human abundance was not significant at the small spatial scale, where both used and available points were far from the areas with higher human abundance. We found bedsite visibility to be a key driver of selection at both spatial scales, with bedsites characterised by significantly lower visibility (denser vegetation) than random available points (Table 2). Bedsites were more likely to be located at distance greater than 50m from roads with vehicle traffic. Finally, bedsite locations were more likely to be selected within forest habitat than open areas (Table 2). We depicted the key scenarios which described bedsite visibility and avoidance of human hotspots and roads in Fig. 5 and Fig. 6. In Fig.5, we showed the highly significant avoidance of human hotspots at the large spatial scale, with bedsites more likely to be located in an area with less than 5 people/ha and at a distance greater than 50m from paved roads. In Fig. 6, we can see for both spatial scales how bedsites were selected in areas with horizontal visibility close to zero and – again – at distance from paved roads greater than 50 meters.

**Fig. 5:**
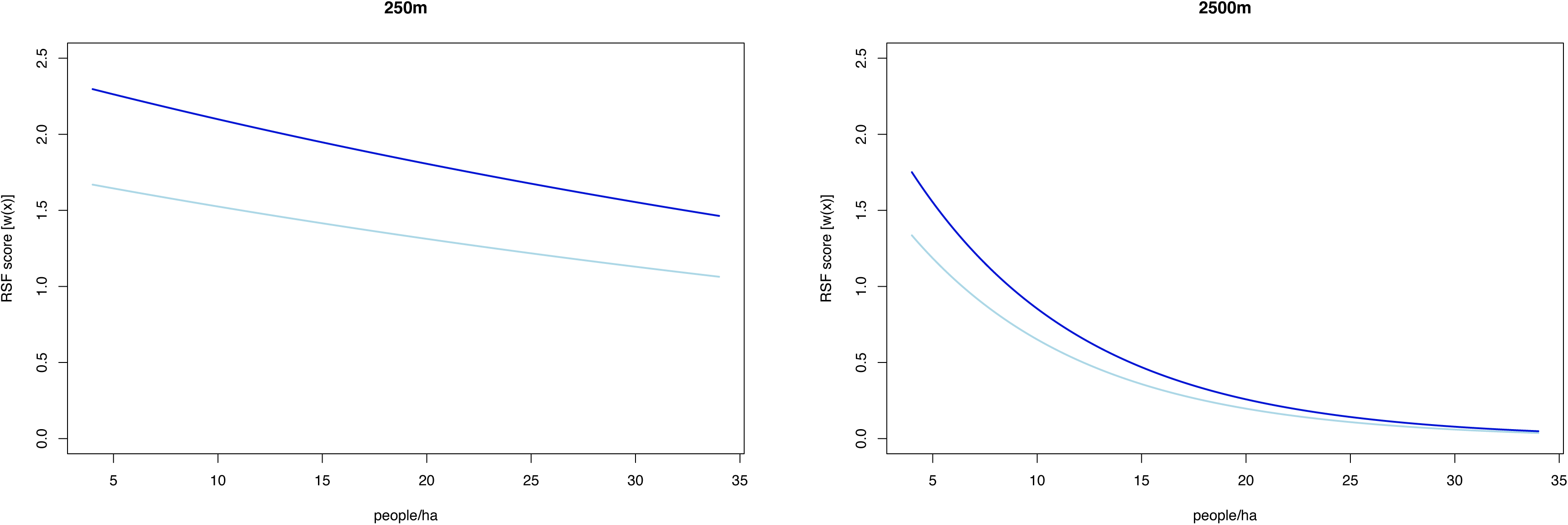
Predicted effect of the number of people per hectare (x-axis, 100×100m) on the relative probability of bedsite selection (y-axis, RSF score [w(x)]) when the distance to the closest road is between 0 and 50 meters from the road (light blue line) or greater than 50 meters from the road (dark blue line). Note the stronger selection for bedsites far from roads, particularly when the number of people per hectare is lower. Also note avoidance of humans was remarkable at the large spatial scale (2500m), and weaker at the small spatial scale (250m), where both bedsites and random available points are both far from human hotspots.

**Fig. 6:**
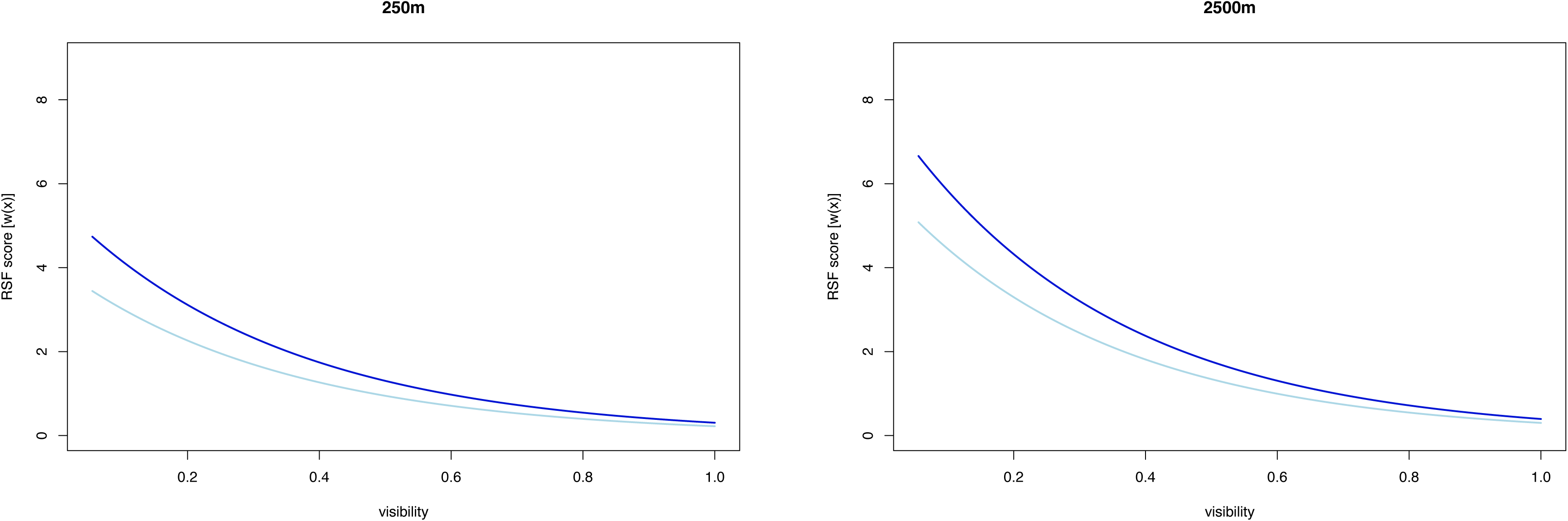
Predicted effect of the bedside visibility (x-axis, ranging from 0% to 100% of a standing fawn visible from 10 meters and eye height of 70cm) on the relative probability of bedsite selection (y-axis, RSF score [w(x)]) when the distance to the closest road is between 0 and 50 meters from the road (light blue line) or greater than 50 meters from the road (dark blue line). Note the stronger selection for bedsites with low visibility and more distant from roads when compared to random available points at both spatial scale.

**Table 2:** Parameters estimated by the GLMMs fitted at two spatial scales (small: 250m; large: 2500m) explaining the relative probability of bedsite selection by fallow deer in the Phoenix Park, Dublin. Significance abbreviations: *** p<0.001, ** p<0.01, * p<0.05, . p<0.1, ns p>0.1, with p < 0.05 corresponding to 95% CI not overlapping zero.

| Scale | Variables | Estimate | Std. Error | Z value | Pr(>\|z\|) | Significance |
| --- | --- | --- | --- | --- | --- | --- |
| **250m** | (Intercept) | -3.39045 | 0.16137 | -21.011 | <2e-16 | *** |
|  | Visibility | -2.30137 | 0.21582 | -10.663 | <2e-16 | *** |
|  | Visibility^2 | 1.29053 | 0.25683 | 5.025 | 5.04e-07 | *** |
|  | Road distance [ >50] | 0.30066 | 0.12802 | 2.349 | 0.0188 | * |
|  | Human abundance | -0.05440 | 0.05392 | -1.009 | 0.3130 | ns |
|  | Human abundance^2 | -0.01307 | 0.05243 | -0.249 | 0.8031 | ns |
|  | Year2019 | -0.97618 | 0.12207 | -7.997 | 1.28e-15 | *** |
|  | Year2020 | -1.33774 | 0.12380 | -10.805 | <2e-16 | *** |
|  | Sex (male) | 0.02158 | 0.10011 | 0.216 | 0.8293 | ns |
|  | Weight | 0.08826 | 0.31559 | 0.280 | 0.7797 | ns |
|  | Weight^2 | -0.11144 | 0.31421 | -0.355 | 0.7228 | ns |
|  | Land cover type (open) | 0.19751 | 0.11180 | 1.767 | 0.0773 | · |
| **2500m** | (Intercept) | -5.77268 | 0.16877 | -34.204 | <2e-16 | *** |
|  | Visibility | -2.37996 | 0.22760 | -10.457 | <2e-16 | *** |
|  | Visibility^2 | 1.29965 | 0.28195 | 4.610 | 4.04e-06 | *** |
|  | Road distance [ >50] | 0.26029 | 0.12131 | 2.146 | 0.0319 | * |
|  | Human abundance | -0.11213 | 0.06320 | -1.774 | 0.0760 | · |
|  | Human abundance^2 | -0.32845 | 0.07786 | -4.219 | 2.46e-05 | *** |
|  | Year2019 | -0.70036 | 0.11676 | -5.998 | 1.99e-09 | *** |
|  | Year2020 | -0.98314 | 0.11871 | -8.282 | <2e-16 | *** |
|  | Sex (male) | 0.03313 | 0.09717 | 0.341 | 0.7331 | ns |
|  | Weight | 0.08524 | 0.31048 | 0.275 | 0.7837 | ns |
|  | Weight^2 | -0.09228 | 0.30750 | -0.300 | 0.7641 | ns |
|  | Land cover type (open) | -0.22119 | 0.10763 | -2.055 | 0.0399 | * |

## Discussion

We found that, when selecting bedsites, fallow deer mothers significantly avoided hotspots of humans on foot (which was also associated with dog presence) along the hiking trail network. Bedsites were located away from paved roads used by vehicular traffic (rarely being located within 50m from a road), mostly within forest habitat, and in dense understory vegetation with low horizontal visibility (nearly zero, from a distance of 10m). For the first time, through the combination of bedsite data and observations of human abundance and dog presence, we described bedsite selection at a high spatial resolution, gathering empirical evidence on the features of bedsites selected by mothers to successfully give birth and protect their fawns during their first weeks of life in a peri-urban setting. This data can be used by urban wildlife managers to identify and protect fawning sites and alleviate human-wildlife conflict during this critical period of the deer annual biological cycle. This is fundamental for maintaining high welfare standards for urban wildlife through the implementation of good management practices and improve human-wildlife coexistence.

Despite both humans on foot and vehicular traffic being highly predictable in the park – cars never leave paved roads and humans on foot rarely leave walking trails – our study showed higher disturbance from humans on foot compared to roads used by vehicular traffic. The first 50m from the nearest paved road was avoided, but bedsites were located at a much higher distance from human hotspots. This may be because vehicles on the road follow a strictly predictable course (vehicles are not allowed off road) compared to people walking/running in the park who could leave the trails (even if they rarely do so) and are often in the company of unleashed dogs. Several studies have shown that wildlife disturbance is highly dependent on the type of human activity (Stankowich, 2008, Naylor et al., 2009, Ciuti et al., 2012). For example, Ciuti et al. (2012) showed that large herbivore behaviour is impacted more by humans on foot than bicycles and horses on trails and roads. Further studies on ungulates have shown human avoidance and behavioural changes when close to locations with high human disturbance levels (Shen-Jin et al., 2007, Whittington et al., 2004, Manor and Saltz, 2004). Mountain Gazelles (*Gazella gazelle*), for example, not only actively avoid humans, but they have also been shown to alter their flight distance when close to hotspots of human presence (Manor and Saltz, 2004). Similar results were found in a study by Shen-Jin et al. (2007); fallow deer were observed to increase vigilance rates when close to locations with higher human density, whereas they increased resting time when further away from humans. Our study adds new empirical evidence on the effects of human presence and activity on ungulate behaviour by reporting novel effects on bedsite selection in a peri-urban setting.

Vehicular disturbance has also been widely documented to impact wildlife behaviour. Noise disturbance created by busy roads, for example, creates a phenomenon called “noise masking”. Noise masking impacts communication between individuals in a wide range of taxa. Many studies have shown severe effects on reproductive success, density and community structure as well as anti-predator and foraging behaviour (Barber et al., 2010, Pelletier, 2014, Papouchis et al., 2001, Bonnot et al., 2013). Most ungulates avoid being close to roads, especially when high-traffic levels are present (Pelletier, 2014, Papouchis et al., 2001). For example, a study by Bonnot et al. (2013) showed that roe deer avoid roads even if it compromises their access to important feeding sites. Similar avoidance behaviour has been shown in our study, although we found that, at this limited spatial scale, fallow deer mothers had to balance a trade-off between finding suitable bedsites and keeping a safe distance from roads, which resulted in avoidance of roads being limited to 50m.

Concealment and safety from predators are the top priorities for ungulate mothers when selecting a bedsite to hide their newborns (Barbknecht et al., 2011). This anti-predator strategy manifests in this study through the selection for low visibility habitats for fawns’ bedsites. Bongi et al. (2008) showed that lactating roe deer females select for habitats with reduced visibility. Likewise, Ciuti et al. (2006) revealed a preference for dense vegetation and low visibility by fallow deer mothers to hide their fawns from predators. We found that this anti-predator behaviour remains intact even within urban areas. Our study demonstrates consistency in the selection for low visibility when choosing fawns’ bedsites at both spatial scales after accounting for the presence of humans on foot (and their dogs) and vehicles on roads. Our findings can therefore inform management decisions when it comes to reducing human pressure on fawning sites within these urban and peri-urban areas. During the fawning season, for instance, hiking trails closer than ∼200m to established fawning sites should be temporarily closed to minimise disturbance, and policies regarding keeping dogs on a leash should be enforced. Understory vegetation within forest habitats as well as grasslands should not be cut (which may be a practice in several urban parks to improve human access): in Phoenix Park, for instance, hiking trails are the only linear features where the grassland is managed, therefore promoting use by park visitors, and making it less likely that they will walk in the high grass closer to fawning areas. Knowing the impact that humans are having on wildlife populations living in urban areas is fundamental to avoid detrimental effects on wildlife welfare and health. The impacts that we have on wildlife can be minimised by understanding the issues present and creating management plans to mitigate them. In this case, wildlife managers working within urban parks and green spaces can incorporate our findings into management plans in order to promote human-deer coexistence during the birthing season.

## Availability of data and materials

The datasets generated and/or analysed during the current study as well as the R markdown script/HTML showing the statistical analysis of this study are available in the ‘*Fawn bedsite selection by a large ungulate living in a peri-urban area*’ repository, https://doi.org/10.5281/zenodo.10886373.

