## Supporting information for "Fawn bedsite selection by a large ungulate living in a peri-urban area"

**Supplementary Table S1**

Collinearity test (Pearson correlation coefficients) of the candidate predictors used to explain bedsite resource selection (presence-available data here labeled as “used”). The sequential colours on the right are associated with the level of correlation between variables with blue indicating a positive correlation (0 to +1) and red a negative correlation (0 to -1). The only variables that were collinear were the predicted number of people (indicating human presence) and likelihood to encounter dogs (indicating canine presence) with a r_p_ > 0.7.

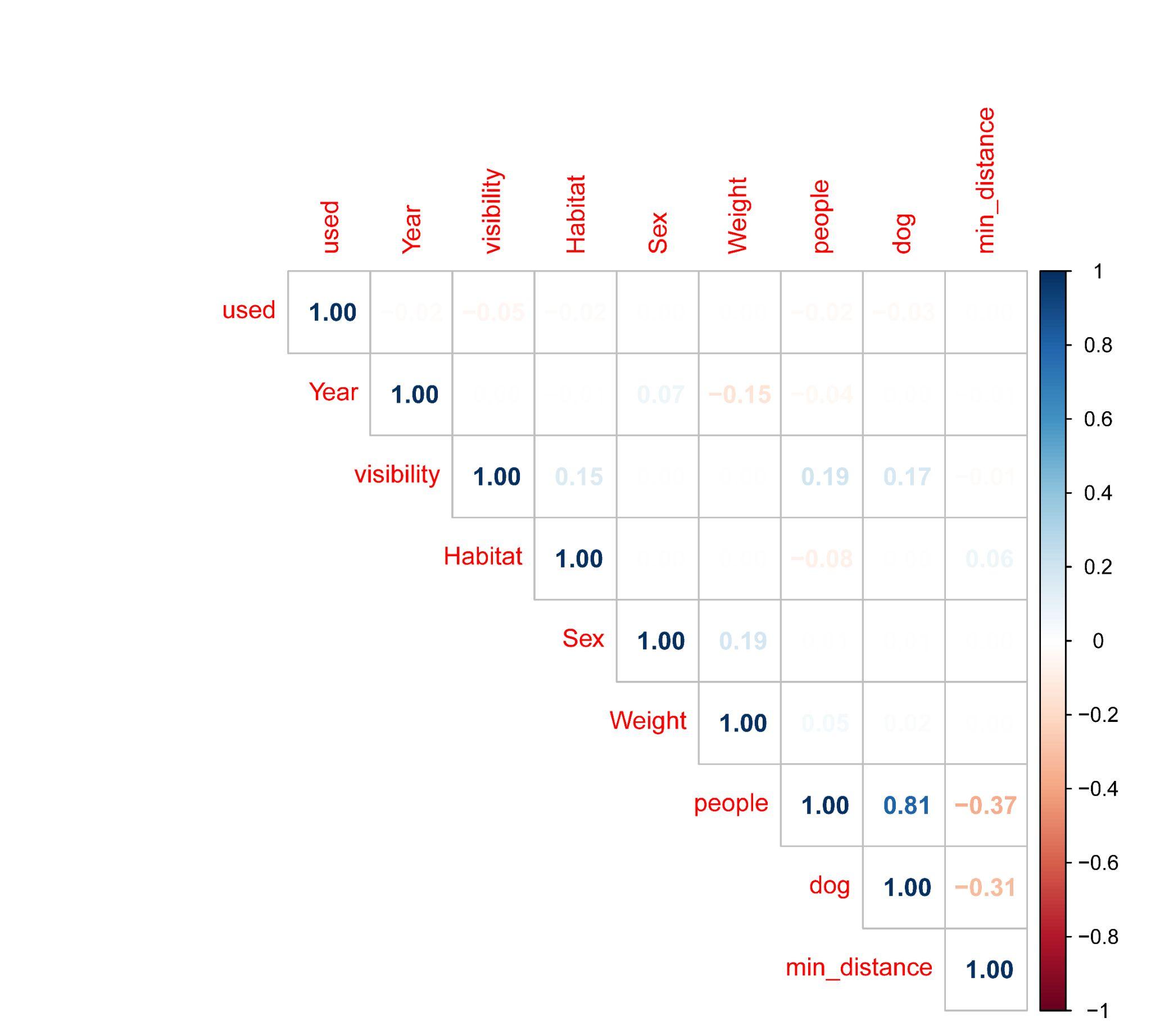

**Supplementary Fig. S1**

Plot of log-likelihood estimates built to estimate the value of lambda (λ) that achieves the optimal transformation for variable describing the number of park visitors (human presence) at the 50 x 50 m resolution.

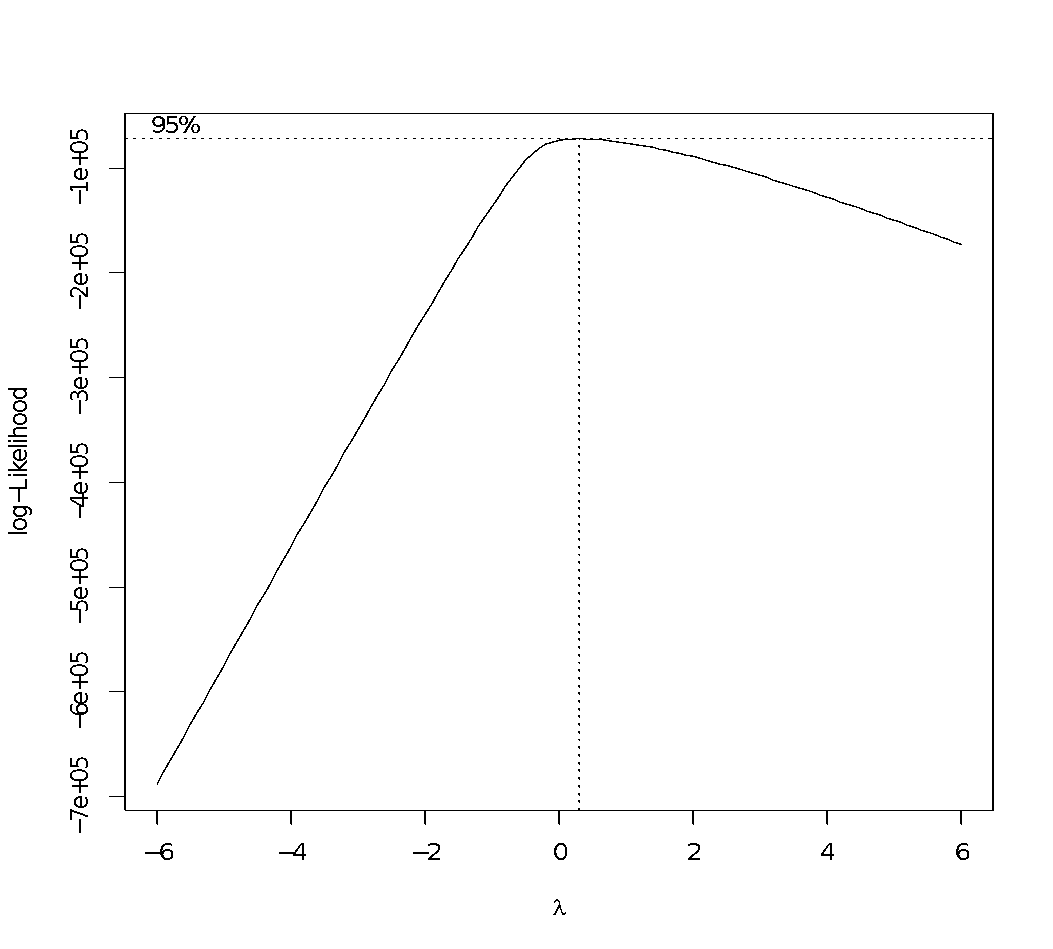

**Supplementary Fig. S2**

Q-Q Plots of the variable describing the number of park visitors (human presence) before (left) and after (right) transformation with Box.Cox.

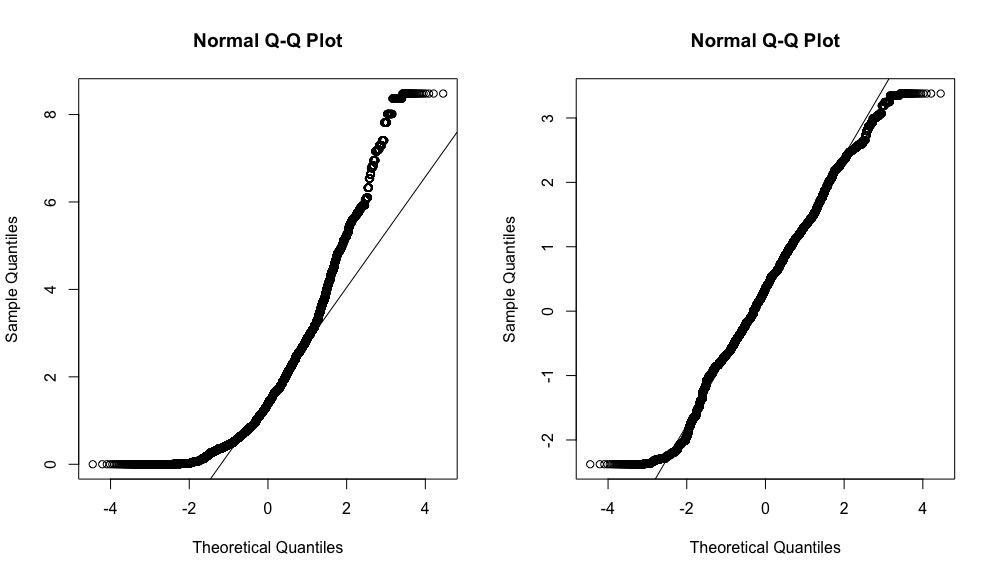

**Supplementary Fig. S3**

AIC values estimated in the sensitivity analyses run to find the best distance threshold (i.e. 50 m) used to categorize the continuous predictor road distance (close: distance < 50m; far: distance >50m).

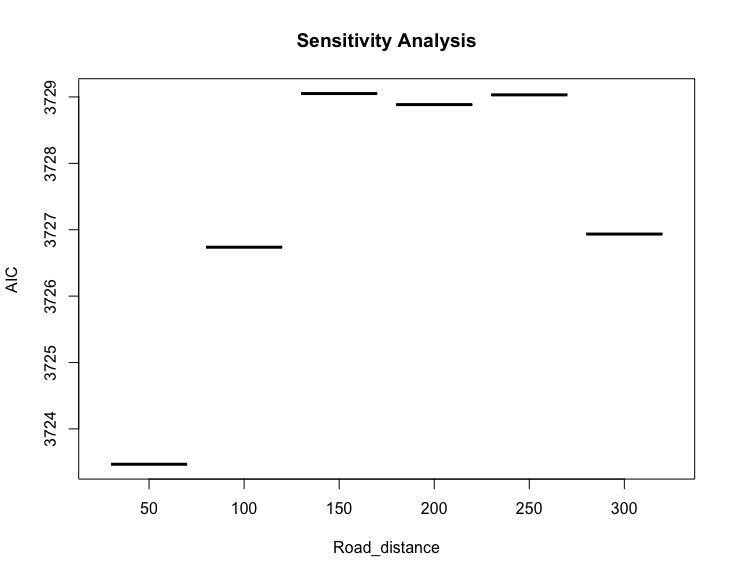

**Supplementary Table S2**

GAM Model 1 (human presence) and Model 2 (dog presence) summary.

**HUMAN MODEL (response: number of park visitor within 50 x 50 m pixel)**

| **Family:** quasipoisson  **Link function:** log | | | | | |
| --- | --- | --- | --- | --- | --- |
| **Formula:**  human presence ~ s(long, lat) + dow | | | | | |
| **Parametric coefficients:** | | | | | |
|  | Estimate | Std. Error | t value | Pr(>\|t\|) |  |
| (Intercept) | -12.1318 | 1.9014 | -6.381 | 2.10e-10 | *** |
| Weekend (opposed to weekday) | 0.6448 | 0.1066 | 6.047 | 1.71e-09 | *** |
| **Approximate significance of smooth terms:** | | | | | |
|  | edf | Ref. df | F | p-value |  |
| s(long, lat) | 28.97 | 29 | 7.406 | <2e-16 | *** |
| **R-sq(adj)** = 0.184 | | | **Deviance explained** = 43.9% | | |
| **GCV** = 2.1399 | | | **Scale est.** = 5.378 | | |
| **n** = 2484 | | | **Significance: 1 (-) 0.1 (.) 0.05(*) 0.001(**) 0 (***)** | | |

**DOG MODEL (response: presence/absence of dogs within 50 x 50 m pixel)**

| **Family:** binomial  **Link function:** logit | | | | | |
| --- | --- | --- | --- | --- | --- |
| **Formula:**  dog presence ~ s(long, lat) + dow | | | | | |
| **Parametric coefficients:** | | | | | |
|  | Estimate | Std. Error | z value | Pr(>\|z\|) |  |
| (Intercept) | -11.2134 | 2.0431 | -5.489 | 4.05e-08 | *** |
| Weekend (opposed to weekday) | 0.2989 | 0.1736 | 1.721 | 0.0852 | . |
| **Approximate significance of smooth terms:** | | | | | |
|  | edf | Ref. df | Chi. sq | p-value |  |
| s(long, lat) | 28.75 | 28.98 | 97.76 | 2.9e-09 | *** |
| **R-sq(adj)** = 0.12 | | | **Deviance explained** = 26.1% | | |
| **UBRE** = -0.61285 | | | **Scale est.** = 1 | | |
| **n** = 2484 | | | **Significance: 1 (-) 0.1 (.) 0.05(*) 0.001(**) 0 (***)** | | |
